# TCCIA: A Comprehensive Resource for Exploring CircRNA in Cancer Immunotherapy

**DOI:** 10.1101/2023.08.24.554049

**Authors:** Shixiang Wang, Yi Xiong, Yihao Zhang, Haitao Wang, Minjun Chen, Jianfeng Li, Peng Luo, Yung-Hung Luo, Markus Hecht, Benjamin Frey, Udo S Gaipl, Xuejun Li, Qi Zhao, Hu Ma, Jian-Guo Zhou

**Affiliations:** Department of Oncology, The Second Affiliated Hospital of Zunyi Medical University, Zunyi, 563000, P. R. China; State Key Laboratory of Oncology in South China, Guangdong Key Laboratory of Nasopharyngeal Carcinoma Diagnosis and Therapy, Guangdong Provincial Clinical Research Center for Cancer, Sun Yat-sen University Cancer Center, Guangzhou 510060, P. R. China; Xiangya School of Medicine, Central South University, Changsha, 410013, P. R. China; Hunan International Scientific and Technological Cooperation Base of Brain Tumor Research, Xiangya Hospital, Central South University, Changsha, 410008, P. R. China; Department of Neurosurgery, Xiangya Hospital, Central South University, Changsha, 410008, P. R. China; Center for Precision Medicine Research and Training, Faculty of Health Sciences, University of Macau, Macau SAR, 999087, P. R. China; State Key Laboratory of Medical Genomics, Shanghai Institute of Hematology, National Research Center for Translational Medicine, Rui-Jin Hospital, Shanghai Jiao Tong University, School of Medicine, Shanghai, 200025, P. R. China; Department of Oncology, Zhujiang Hospital, Southern Medical University, Guangzhou, 510091, P. R. China; Department of Chest Medicine, Taipei Veterans General Hospital, Taipei, 11217, Taiwan (R.O.C.); School of Medicine, College of Medicine, National Yang Ming Chiao Tung University, Taipei, 112304, Taiwan (R.O.C.); Department of Radiotherapy and Radiation Oncology, Saarland University Medical Center, Homburg, 66421, Germany; Translational Radiobiology, Department of Radiation Oncology, Universitätsklinikum Erlangen, Friedrich-Alexander-Universität Erlangen-Nürnberg, Erlangen, 91054, Germany

**Keywords:** Circular RNA, Database, Checkpoint immunotherapy, Biomarker

## Abstract

**Background:** Immunotherapies targeting immune checkpoints have gained increasing attention in cancer treatment, emphasizing the need for predictive biomarkers. Circular RNAs (circRNAs) have emerged as critical regulators of tumor immunity, particularly in the PD-1/PD-L1 pathway, and have shown potential in predicting immunotherapy efficacy. Yet, the detailed roles of circRNAs in cancer immunotherapy are not fully understood. While existing databases focus on either circRNA profiles or immunotherapy cohorts, there is currently no platform that enables the exploration of the intricate interplay between circRNAs and anti-tumor immunotherapy. A comprehensive resource combining circRNA profiles, immunotherapy responses, and clinical outcomes is essential to advance our understanding of circRNA-mediated tumor-immune interactions and to develop effective biomarkers.

**Methods:** To address these gaps, we constructed the Cancer CircRNA Immunome Atlas (TCCIA), the first database that combines circRNA profiles, immunotherapy response data, and clinical outcomes across multi-cancer types. The construction of TCCIA involved applying standardized preprocessing to the raw sequencing FASTQ files, characterizing circRNA profiles using an ensemble approach based on four established circRNA detection tools, analyzing tumor immunophenotypes, and compiling immunotherapy response data from diverse cohorts treated with immune-checkpoint blockades (ICBs).

**Results:** TCCIA encompasses over 4,000 clinical samples obtained from 25 cohorts treated with ICBs along with other treatment modalities. The database provides researchers and clinicians with a cloud-based platform that enables interactive exploration of circRNA data in the context of ICB. The platform offers a range of analytical tools, including browse of identified circRNAs, visualization of circRNA abundance and correlation, association analysis between circRNAs and clinical variables, assessment of the tumor immune microenvironment, exploration of tumor molecular signatures, evaluation of treatment response or prognosis, and identification of altered circRNAs in immunotherapy-sensitive and resistant tumors. To illustrate the utility of TCCIA, we showcase two examples, including circTMTC3 and circMGA, by employing analysis of large-scale melanoma and bladder cancer cohorts, which unveil distinct impacts and clinical implications of different circRNA expression in cancer immunotherapy.

**Conclusions:** TCCIA represents a significant advancement over existing resources, providing a comprehensive platform to investigate the role of circRNAs in immuno-oncology.

*What is already known on this topic:* Prior knowledge indicated that circRNAs are involved in tumor immunity and have potential as predictive biomarkers for immunotherapy efficacy. However, there lacked a comprehensive database that integrated circRNA profiles and immunotherapy response data, necessitating this study.

*What this study adds:* This study introduces TCCIA, a database that combines circRNA profiles, immunotherapy response data, and clinical outcomes. It provides a diverse collection of clinical samples and an interactive platform, enabling in-depth exploration of circRNAs in the context of checkpoint-blockade immunotherapy.

*How this study might affect research, practice or policy:* The findings of this study offer valuable insights into the roles of circRNAs in tumor-immune interactions and provide a resource for researchers and clinicians in the field of immune-oncology. TCCIA has the potential to guide personalized immunotherapeutic strategies and contribute to future research, clinical practice, and policy decisions in checkpoint-blockade immunotherapy and biomarker development.

## Introduction

Immunotherapy has revolutionized the treatment of cancer over the past decade, emerging as a groundbreaking approach that harnesses the patient’s own immune system to fight cancer. Therapies like immune checkpoint blockades (ICBs), chimeric antigen receptor (CAR) T-cell therapy, and therapeutic vaccines aim to reinvigorate anti-tumor immunity against malignant cells [1–3]. ICBs targeting programmed cell death protein-1 (PD-1), PD-1 ligand (PD-L1), and cytotoxic T-lymphocyte-associated protein 4 (CTLA-4) in particular have demonstrated remarkable clinical efficacy across diverse cancer types, highlighting immunotherapy’s potential for a durable and curative response [4]. Adoptive cell transfer using engineered T cells expressing CARs has also shown great promise for blood cancers [5]. However, significant challenges remain in extending immunotherapies to larger patient populations and solid tumors. Heterogeneity in response is a major limitation – while some patients achieve long-term remission, others exhibit intrinsic resistance or relapse after an initial response [6]. This variable efficacy likely stems from immunosuppressive mechanisms within the tumor microenvironment (TME) that enable cancer cells to evade immune attack [7]. Elucidating the complex cellular and molecular interactions underlying immunotherapy resistance will be critical to unlock the full potential of immune-based cancer treatments. Reliable predictive biomarkers are also imperative to guide patient selection and combination immunotherapies tailored to each patient’s TME [8].

Circular RNAs (circRNAs) have recently emerged as fascinating non-coding RNA regulators with unique covalently closed loop structures. Initially considered splicing byproducts, circRNAs are now recognized as important gene expression modulators with diverse functions [9,10]. In cancer, circRNAs have been implicated in proliferation, metastasis, and malignancy hallmarks [11]. Moreover, circRNAs are now recognized as critical regulators and potential biomarker of tumor immunity and immunotherapy response [7,12,13]. Accumulating evidence indicates circRNAs modulate TME and immunotherapy outcomes through various mechanisms in cancers like lung cancer, melanoma, colorectal cancer, and pancreatic cancer [14,15]. For example, circRNAs such as circFGFR1, circ-CPA4, and circ_0000284 facilitate immune evasion by modulating PD-L1 via sponging tumor-suppressive microRNAs [16–18]. Additionally, circRNAs including hsa_circ_0000190 [19], circ_0020710 [20], CDR1-AS [16], and circ-UBAP2 [21] upregulate immune checkpoint proteins like PD-L1, CTLA-4 and PD-1, hampering T cell function and promoting immune evasion. Furthermore, cancer cell-derived circRNAs can reprogram intratumoral immune cells via exosomal transfer or cytokine signaling, thereby impacting facets like angiogenesis that affect immunotherapy efficacy [22–24]. CircRNAs influence various aspects of the TME, including vascularization [25], metabolism [26], hypoxia [27], macrophage polarization [28], natural killer cell cytotoxicity [17], and T cell exhaustion/apoptosis [29]. These factors can impede the efficacy of immunotherapy [15,30]. Dysregulation of circRNAs promotes immune destruction evasion and reduced immunotherapy efficacy. CircRNAs employ diverse regulatory mechanisms—from sponging miRNAs and proteins to scaffolding proteins and translating peptides [31]. While many circRNAs originate in tumors, others come from stromal and immune cells, underscoring complex multicellular regulation [32]. Exploring circRNA networks will be critical to unraveling this intricate cancer-immunity interplay. With emerging roles in tumor immunity, prognostic potential, and biomarker utility, circRNAs represent a promising new frontier in cancer immunotherapy.

Despite growing interest in circRNAs and their potential relevance in cancer immunotherapy, a comprehensive understanding of their precise functions and clinical implications remains incomplete. Existing databases have limitations in either profiling circRNAs, such as riboCIRC [33], CSCD [34] and CircNet [35] offering circRNA profiles across tissues or cancers, or curating immunotherapy cohorts, like ICBatlas [36] and TCIA [37] compiling immune infiltration and immunotherapy data across tumor types. Crucially, no resources systematically integrate comprehensive circRNA expression with multi-omics datasets including immune cell fractions, ICB types, and clinical outcomes for systematic exploration of the circRNA-immunotherapy interplay.

To address this unmet need, we developed the first-of-its-kind database, The Cancer CircRNA Immunome Atlas (TCCIA), a comprehensive database that integrates circRNA profiles, immunotherapy response data, and clinical outcomes for multiple cancer types, with the objective of providing a valuable resource for systematic exploration of the circRNA-immune axis, advancing our understanding of their functions and to facilitates discovery of potential biomarkers, therapeutic targets and clinical implications in cancer immunotherapy.

## Results

### Integrating circular RNAs in cancer immunotherapy

The development of the TCCIA database encompassed a comprehensive process involving data collection, preprocessing, and integration (Figure 1). In terms of data collection, we carefully curated research articles detailing cohorts treated with ICBs, utilizing the PubMed database (https://www.ncbi.nlm.nih.gov/pubmed/) for selection. We acquired raw RNA-seq datasets from several genome sequence archive repositories, including dbGaP, EGA, EMBL-EBI and GSA. We applied standardized preprocessing techniques to identify circRNAs, quantify TME/gene signatures, and incorporate clinical annotations and outcomes. This approach was aimed at improving the consistency and comparability of data across different datasets. Notably, TCCIA addresses a crucial gap by providing a unified platform with multiple tools that facilitates the exploration of circRNAs’ impact on immunotherapy outcomes (Figure 1). This sets it apart from many existing databases [33–47] that predominantly focus on circRNA profiles, circRNA annotations, or immunotherapy cohorts (Table 1).

**Figure 1.**
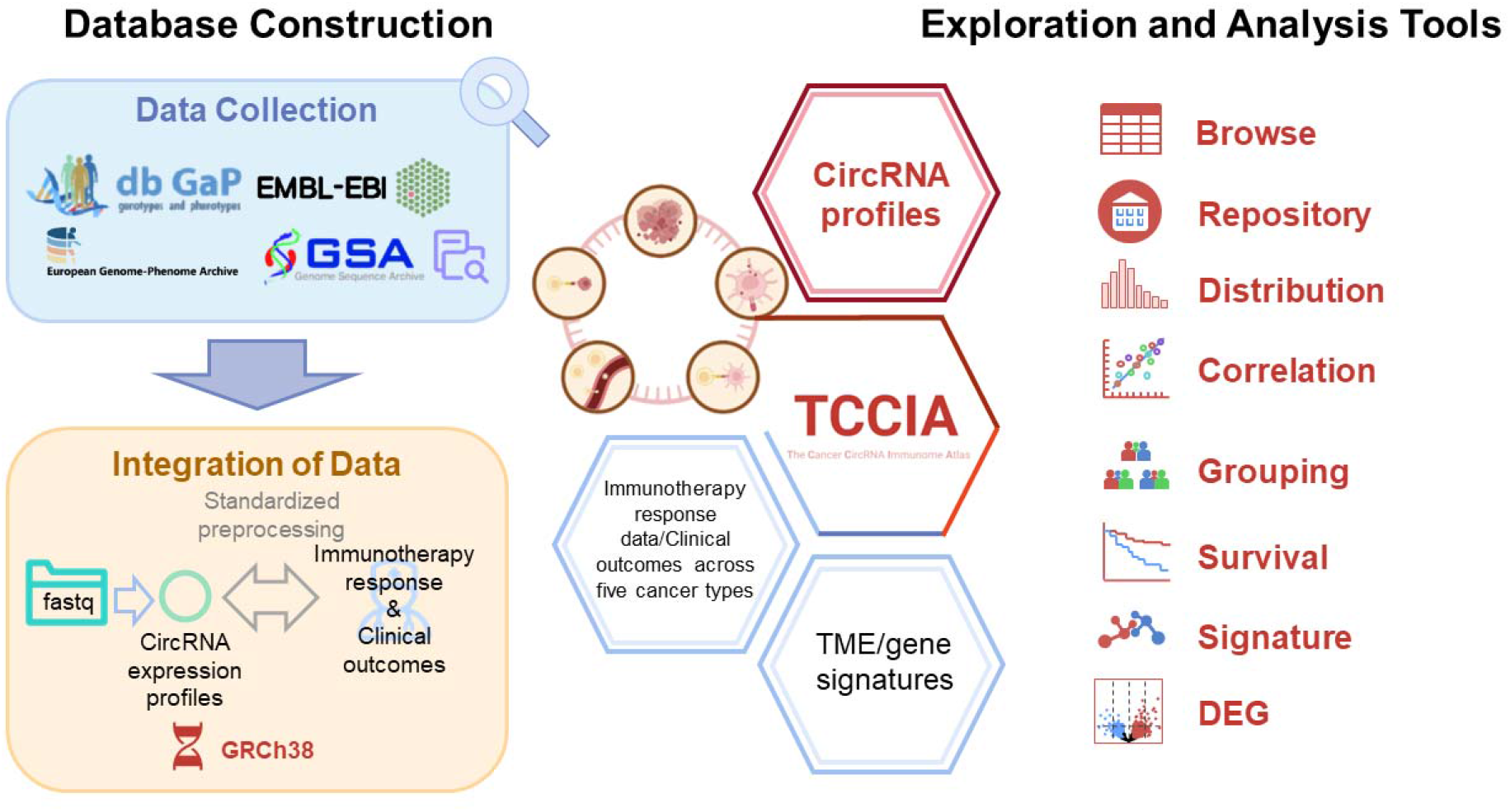
Overview of TCCIA.

**Table 1.** Comparison between TCCIA and other similar database resources [33–47].

| Type | Name | Publication (Year) | Last update | Support species | Sample types (N) | Number of samples | Data sources | Characteristics |
| --- | --- | --- | --- | --- | --- | --- | --- | --- |
| Mixed | TCCIA | Submitted | 2023 | Homo sapiens | cancer (10) | > 4000 cancer samples | dbGaP, EMBL-EBI, EGA, GSA | Characterization of circRNA expression profiles within and across checkpoint blockade immunotherapy clinical datasets; Comparison of circRNA between clinical phenotypes (including response vs. nonresponse in pre-treatment or on-treatment); circRNA based survival analyses; the association of circRNA with 255 cancer signatures and 8 types of TME metrics; analyzing multiple circRNAs simultaneously |
| CircRNA related | CSCD2 | NAR Database Issue (2022) | 2022 | Homo sapiens | cancer (23) | 825 tissues + 288 cell lines | ENCODE, SRA | Prediction of potential miRNA–circRNA and RBP–circRNA interactions; prediction of potential circRNA full-length and open reading frame sequence |
|  | circMine | NAR Database Issue (2022) | 2022 | Homo sapiens | disease (87 ) | 1107 samples | GEO | Provide 13 online analytical functions to comprehensively investigate circRNAs, including search, browse, differential expression analysis, co-expression analysis, circRNA-miRNA prediction, circRNA IRES prediction, ribo-circRNA location, etc. |
|  | CircR2Disease2 | Genomics, Proteomics & Bioinformatics (2021) | 2022 | 5 species | disease (313) | 2449 studies | PubMed | Investigation of dysregulated circRNAs and their posttranscriptional regulatory function |
|  | CircNet2 | NAR Database Issue (2021) | 2021 | Homo sapiens | cancer (37) | 2732 cancer samples | TCGA, GEO, CircAtlas, MiOncoCirc | Characterization of tissue-specific circRNA expression profiles and circRNA-miRNA-gene regulatory network |
|  | riboCIRC | Genome Biology (2021) | 2021 | 21 species | tissue/cell line (314) | 1970 samples | GEO | Exploration of computationally predicted ribosome-associated circRNAs, experimentally verified translated circRNAs, and cross-species conserved ribo-circRNAs |
|  | CircAtlas2 | Genome Biology (2020) | 2020 | 7 species | tissue (20) | 1070 samples | SRA, NGDC, GeneBank | Comprehensive profiling of circRNAs, including their searching, orthologous, converter, expression, sequences, translations, and functions |
|  | CircInteractome | RNA Biology (2016) | 2020 | Homo sapiens | tissue/cell line (34) | 34 samples | circBase | Exploration of the interactions between miRNAs and circRNAs with 109 RBPs |
|  | TransCirc | NAR Database Issue (2020) | 2020 | Homo sapiens | tissues (17) | 17 tissues | CircAtlas | Search and browse six types of evidence supporting the translation potential of circRNAs |
|  | circRic | Genome Medicine (2019) | 2019 | Homo sapiens | cancer (22) | 935 cancer celllines | CCLE | Characterization of circRNA expression profiles; analysis of the circRNA biogenesis regulators, the effect of circRNAs on drug response, the association of circRNAs with mRNAs, proteins, mutations, and RBPs and miRNA binding sites. |
|  | CIRCpedia2 | Genomics, Proteomics & Bioinformatics (2018) | 2018 | 6 species | tissue (13) | 185 samples | GEO, ENCODE, EMBL-EBI | Search, browse and download CIRCexplorer2 identified circRNAs; simple visualization of circRNA expression |
|  | circBase | RNA (2014) | 2014 | 6 species | tissues/cell line (77) | 77 samples | GEO, ENCODE, EMBL-EBI | Public (evidence supporting) circRNA search; annotation preview and export |
| Immunotherapy related | TISMO | NAR Database Issue (2022) | 2022 | Mouse | cancer (19) | 1518 mouse samples | GEO and In-house data | Summary data browser; Boxplots comparing expression of a gene, pathway, or immune cell type either in vivo or in vitro samples across treatment and response conditions |
|  | ICBAtlas | Cancer Immunol Res (2022) | 2022 | Homo sapiens | cancer (9) | 1515 cancer samples | GEO,ArrayExpress, TCGA, dbGaP | Gene-, pathway-, and immune cell-level search, summary, and visualization; Comparison between response vs. nonresponse, pre-treatment vs. on-treatment of checkpoint blockade immunotherapy; Provide a response score system |
|  | ImmuCellAI | Advanced Science (2020) | 2020 | Homo sapiens | cancer (37) | NA | GEO, TCGA, dbGaP | Estimation of the abundance of 24 immune cells; survival analysis |
|  | TCIA | Cell Reports (2017) | 2017 | Homo sapiens | cancer (20) | 9562 cancer samples | TCGA and two immunotherapy studies | Advanced data selection; gene expression analysis; cell type abundance analysis; GSEA; survival analysis; tumor heterogeneity summary; neoantigen summary and analysis |

### Data summary of TCCIA

In this study, a comprehensive compilation was made, involving approximately 4000 clinical patient samples from 25 immune-checkpoint blockade (ICB) cohorts [48–70] with raw RNA-seq datasets published between 2012 and 2023, encompassing 10 distinct cancer types (Figure 2A, B and Supplementary Table 1). We employed an ensemble strategy that combined the outcomes of four different circRNA detection tools: CIRCexplorer2 [71], CIRIquant [72], find_circ [73], and circRNA_finder [74]. This approach is akin to the one utilized in the study conducted by Dong *et al*. [13] (refer to the Methods section for more details). In total, we detected 793,316 circRNAs using at least two circRNA detection methods. By applying the ensemble strategy, we successfully identified and included a total of 555,098 unique circRNAs in the TCCIA database (Figure S1). A significant proportion of these circRNAs (23.2% to CIRCpedia [44] and 40.3% to circAltas [40]) could be confidently mapped to well-known circRNA databases. Among included cancer types, Bladder urothelial carcinoma (BLCA) exhibited the highest level of circRNA abundance, encompassing 141,085 circRNAs (representing 25.4% of the total). Conversely, Sarcoma (SARC) demonstrated the lowest level of circRNA abundance, with only 4109 circRNAs, constituting a mere 0.7% of the total (Figure 2B). The average lengths of these circRNAs demonstrated notable consistency across various cancer types, ranging from 11,176.81 to 19,970.70, as well as within individual cohorts, ranging from 8,899.24 to 21,357.33 (Figure 2C). Intriguingly, Head and neck squamous cell carcinoma (HNSC) exhibited the highest mean back-splice junction (BSJ) reads at 5.10, whereas BLCA had the lowest mean at 0.07 (Figure 2C). A comprehensive analysis encompassing all the circRNAs from the sampled datasets was visualized using t-SNE (Figure 2D). This visualization revealed discernible circRNA clustering patterns specific to various cancer types, highlighting the nuanced circRNA heterogeneity within human cancers and emphasizing the need for independent circRNA analysis considerations.

**Figure 2.**
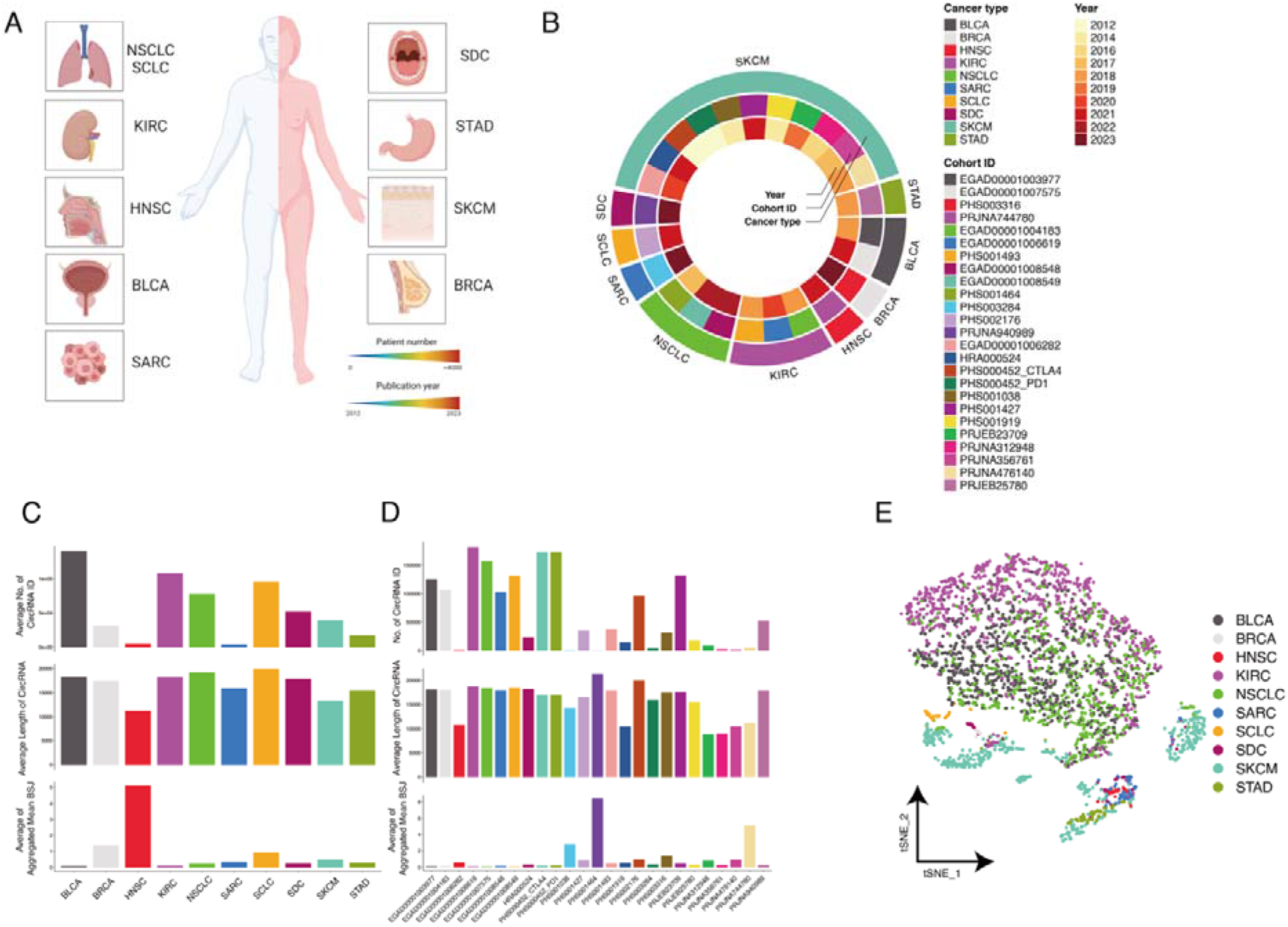
Content of TCCIA. (**A**) Inclusion of cancer types, cohort and publication years in this study. **Abbr.** BLCA, Bladder urothelial carcinoma; HNSC, Head and neck squamous cell carcinoma; KIRC, Kidney renal clear cell carcinoma; NSCLC, Non-small lung cancer; SCLC, Small lung cancer; SKCM, Skin cutaneous melanoma; SGC, Salivary gland carcinoma; BRCA, Breast cancer; SARC, Sarcoma; STAD, Stomach Adenocarcinoma. (**B-C**) The number of detected circRNAs, length of circRNAs, mean (back-splicing junction count) BSJ and mean (counts per million) CPM in different cancer types (**B**) or cohort (**C**). (**D**) UMAP plot of all samples, colored by cancer types.

### Web functionality of TCCIA

TCCIA offers a diverse array of exploratory and advanced analytical tools, providing a comprehensive platform for researchers to uncover intricate connections and gain valuable insights (Figure 1). The modules are thoughtfully introduced, organized, and accessible across nine web pages: "Home", "Browse", "Repository", "Cohort-centred Analysis", "Molecule-centred Analysis", "Signature", "DE circRNAs”, "Setting", and "Help" These functionalities enable users to explore circRNA information, abundance, correlations with clinical variables, the tumor immune microenvironment, molecular signatures, treatment responses, and prognosis predictions. Researchers can also identify circRNAs associated with both immunotherapy-sensitive and resistant tumor scenarios. The well-established exploration and analysis workflow within the TCCIA framework is illustrated in Figure 3, outlining the typical path that researchers follow when engaging with the platform. A more comprehensive explanation of each fundamental module is provided below.

**Figure 3.**
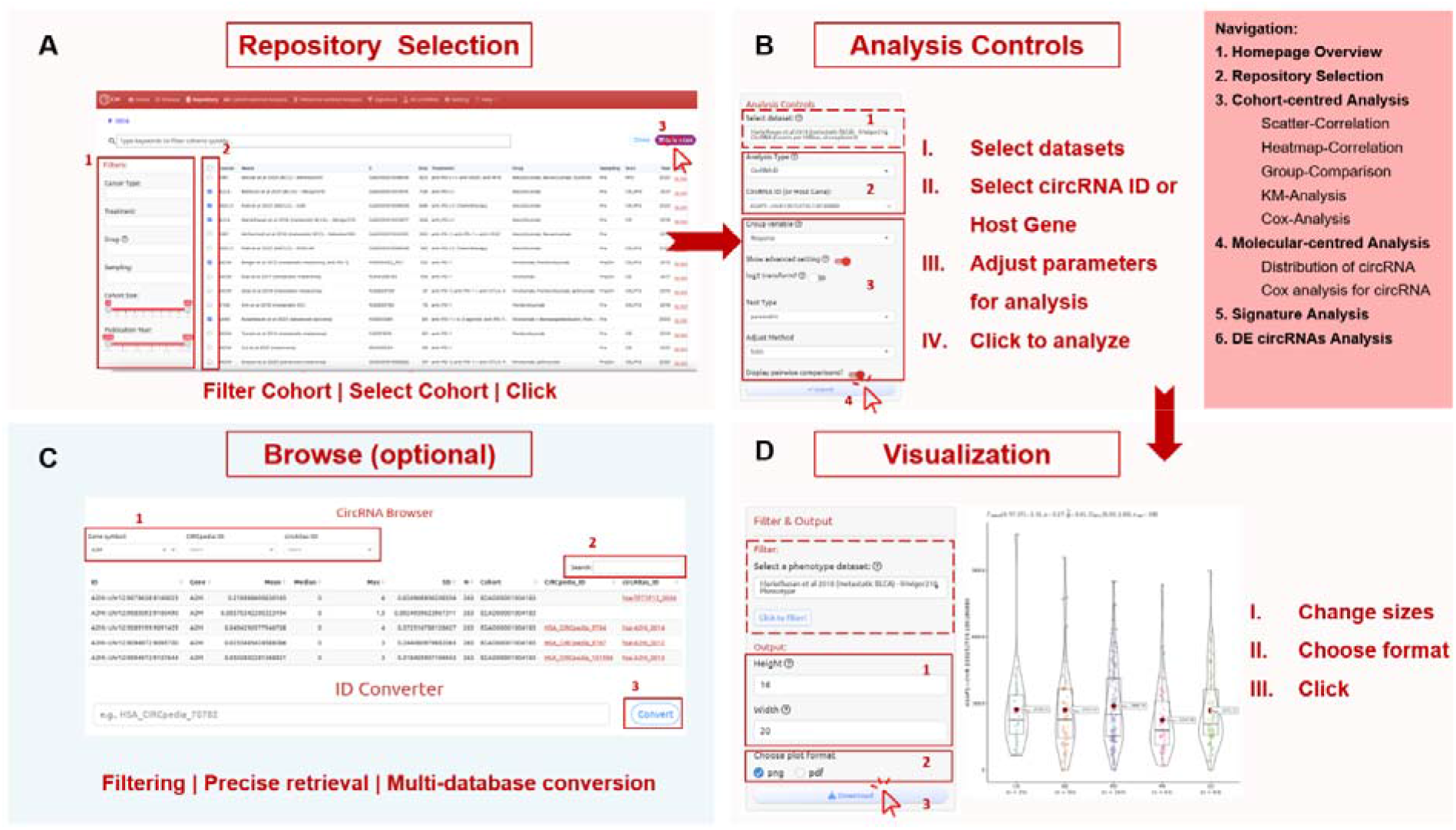
A standard exploration and analysis pipeline within TCCIA. **(A)** Users need to navigate to the Repository page and select the clinical cancer dataset(s) of their interest. On the leftmost panel, users can filter datasets based on various parameters. **(B)** Users can choose an appropriate analysis strategy according to their needs (core analysis steps are listed on the rightmost panel). The analysis typically consists of four steps: I. Select the dataset(s). II. Choose the CircRNA ID or Host Gene of interest. III. Adjust various parameters such as Test Type, Color Selection, etc. IV. Click the Submit button to obtain the analysis and visualization results. **(C)** Users can browse various information about all CircRNAs included in TCCIA, and at the same time, they can use the ID Converter to transform their interested CircRNAs into other CircRNA Database IDs. **(D)** Users can use the provided options to save result images in PDF or PNG format, with the desired dimensions.

#### CircRNA Browser

TCCIA’s circRNA browser offers a distinctive and valuable resource for the exploration and retrieval of identified circRNAs within cancer cohorts. The browser provides essential information such as gene symbols, genomic locations (based on the hg38 genome build), cohort origins, statistical metrics reflecting circRNA expression distribution (including mean, median, maximum, standard deviation, and sample size - N), and cross-references to identifiers from CIRCpedia and circAtlas. Users can conveniently pinpoint circRNAs of interest by applying filters based on gene symbols, CIRCpedia identifiers, and/or circAtlas identifiers.

#### Cohort Selection and Data Access

The TCCIA interface offers an intuitive approach for cohort selection and data access. The Repository Page serves as a gateway, enabling users to filter datasets based on crucial parameters such as cancer type, treatment modalities, drugs administered, and cohort sizes. Essential details pertaining to each dataset are presented in a comprehensive cohort table, facilitating informed decision-making regarding cohort selection.

#### Cohort/Molecule-Centered Analysis Modules

At the heart of TCCIA’s capabilities lie the cohort-centered analysis modules and molecule-centered analysis modules (for analyzing circRNAs across multiple cohorts), providing a profound lens into circRNA dynamics within specific immunotherapy cohorts. These modules encompass:

1. *Scatter-Correlation and Heatmap-Correlation*: Researchers gain insights into circRNA correlations through scatter plots and heatmaps. These visualizations are pivotal in elucidating potential connections between circRNAs and other variables within the chosen cohort.
2. *Group-Comparison (including simplified and comprehensive versions)*: TCCIA facilitates nuanced analysis of numeric differences in circRNA expression across multiple groups within a cohort. The dual modes of simplified and comprehensive group comparison empower researchers to unravel intricate circRNA expression patterns.
3. *Kaplan-Meier(KM)-Analysis and Cox-Analysis*: Survival analysis is made accessible through the KM-Analysis module, which generates Kaplan Meier survival curves among distinct variable groups. Additionally, the Cox-Analysis module allows for an in-depth examination of survival outcomes of any circRNA expression, opening avenues to prognostic evaluations.

#### Signature and DEG Analysis

TCCIA introduces dedicated modules for signature analysis and differential expression circRNAs (DEG) assessment. The Signature Page facilitates the investigation of associations between circRNAs and tumor microenvironment metrics using eight prominent deconvolution methods. It also allows for the examination of connections between circRNAs and 255 cancer signatures categorized into three distinct groups: TME-associated, tumor-metabolism, and tumor-intrinsic signatures. These analyses encompass a wide range of cohorts, ensuring comprehensive exploration of these relationships. The DEG Page empowers researchers to pinpoint differentially expressed circRNAs between patients who respond and those who do not respond to immunotherapy, thus unraveling the intricate web of circRNA involvement in treatment outcomes.

#### User Customized Configurations

Global settings within TCCIA add a layer of refinement to the user experience, allowing for customized exploration. These settings grant users’ control over data access and enable tailoring analyses to align with their specific research objectives. For example, by default, the platform prioritizes immunotherapy-related sample data by filtering out samples without checkpoint immunotherapy treatment, streamlining analyses for coherent research goals. Besides, TCCIA supports gene signature in a nutshell, i.e., the addition, subtraction, multiplication, and division are support for aggregating circRNAs of interest as a whole. As users become acclimated to the platform, customization options foster enhanced flexibility, enabling researchers to uncover novel insights.

In essence, the web functionality of TCCIA embodies an advanced and user-centric avenue for investigating the complex roles of circRNAs in cancer immunotherapy. The integration of diverse analysis modules, coupled with a cohort-centered approach and adaptable settings, positions TCCIA as an indispensable tool for advancing our comprehension of circRNA-mediated immune responses and guiding the formulation of personalized immunotherapeutic strategies. This interactive platform stands poised to reshape the landscape of circRNA-immunotherapy research.

### Case studies validating and extending data of two reported CircRNAs

ICB therapies targeting PD-1 and CTLA-4 have significantly transformed the field of oncology, particularly in the treatment of metastatic melanoma. However, it is important to note that only a limited number of melanoma patients experience positive outcomes from these immunotherapies. Consequently, there is a pressing need to identify predictive biomarkers that can guide precision oncology.

In the study conducted by Dong *et al*. [13], it was observed in metastatic melanoma patients from the Gide *et al*. cohort [54] and independently validated in metastatic melanoma patients from the Berger *et al*. cohort [63], who exhibited high expression of circTMTC3, experienced poorer survival outcomes and demonstrated reduced treatment responsiveness compared to those with low circTMTC3 expression. To illustrate the functionality of the TCCIA, here, we performed a reproducible analysis on the same cohorts (Figure 4).

**Figure 4.**
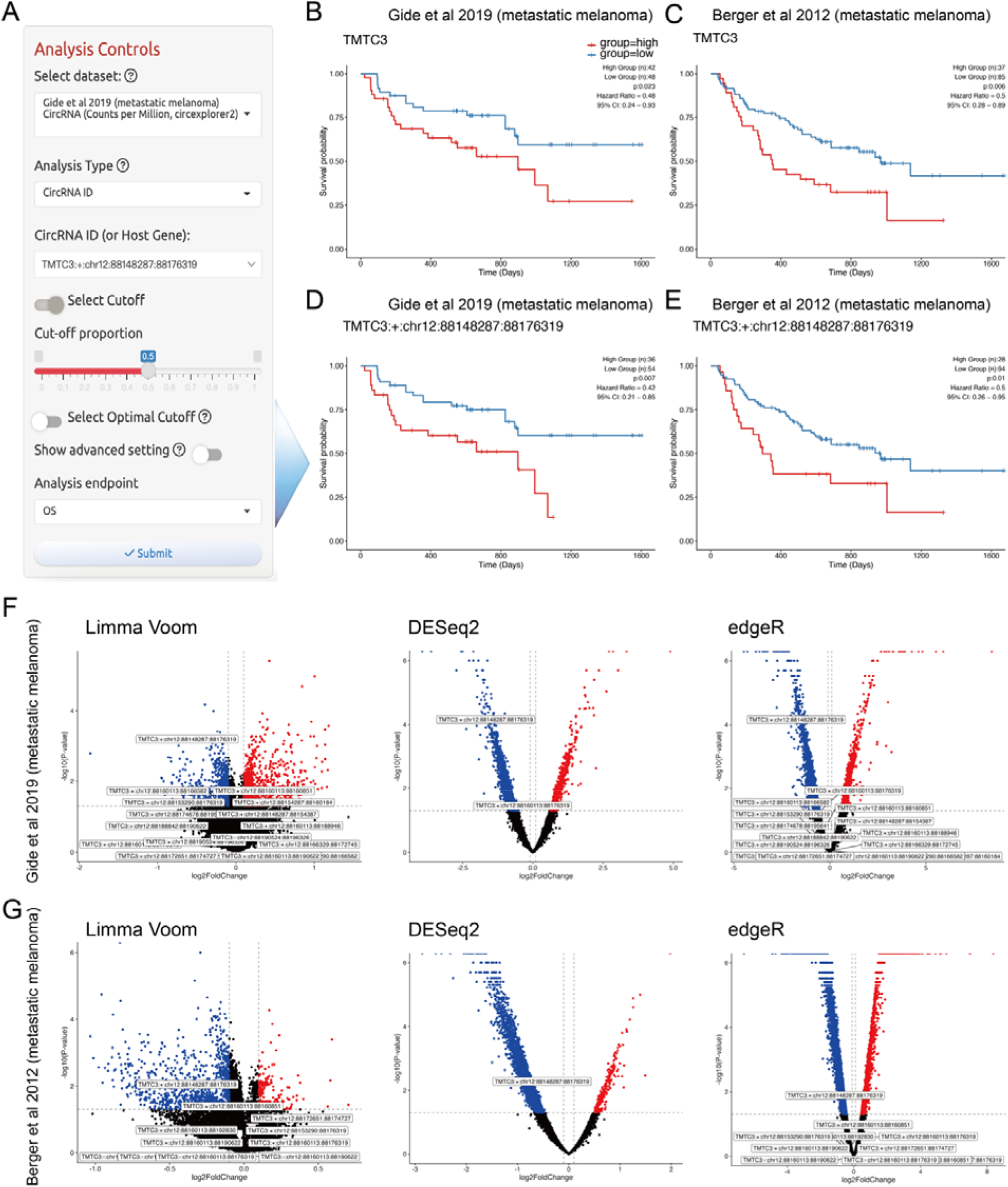
circTMTC3 predicts poor survival and non-response to checkpoint immunotherapy in the melanoma cohorts of Gide *et al*. and Berger *et al*. (**A**) Analysis panel for analyzing and visualizing associations between circRNAs and patient survival. Here, we present the data and analysis settings used to generate the plot shown in (**D**). (**B-C**) CircTMTC3 predicts poor survival in (**B**) Gide *et al*. cohort and (**C**) Berger *et al*. cohort. (**D-E**) A circRNA isoform of circTMTC3, TMTC3:+:chr12:88148287:88176319, predicts poor survival in (**D**) Gide *et al*. cohort and (**E**) Berger *et al*. cohort. Red, group of high circRNA expression. Blue, group of low circRNA expression. (**F-G**) Volcano plots showing differential expressed circRNAs between checkpoint immunotherapy response and nonresponse patients using the approaches Limma Voom, DESeq2, and edgeR in (**F**) Gide *et al*. cohort and (**G**) Berger *et al*. cohort.

TCCIA provides an overview of the analysis panel used to assess the association between a circRNA and patient survival (Figure 4). Here, using TCCIA clearly demonstrate that high levels of circTMTC3 are predictive of poor overall survival in the Gide *et al*. cohort (HR_low vs high_=0.48, 95% CI: 0.24-0.93, *P*=0.023) and the Berger *et al*. cohort (HR_low vs high_=0.5, 95% CI: 0.28-0.89, *P*=0.006) (Figure 4B-C). Upon closer examination of the different isoforms of circRNAs derived from TMTC3, it was found that only TMTC3:+:chr12:88148287:88176319 was abundant and played a significant role in predicting unfavorable survival outcomes in the Gide *et al*. cohort (Figure 4D, HR_low vs high_=0.42, 95% CI: 0.21-0.85, *P*=0.007) and the Berger *et al*. cohort (Figure 4E, HR_low vs high_=0.5, 95% CI: 0.26-0.95, *P*=0.01). Furthermore, we utilized three commonly employed methods to evaluate the differential expression of circRNAs between checkpoint immunotherapy responders and non-responders. Consistent with the survival outcome, we observed that TMTC3:+:chr12:88148287:88176319 was the only isoform of circTMTC3 to exhibit significant upregulation in non-responding patients from both metastatic melanoma cohorts (Figure 4F-G). Mechanistically, Dong *et al*. [13] demonstrated that the overexpression of circTMTC3 could elevate PD-L1 expression through the miR-142-5p/PD-L1 axis, thereby reducing T cell activity and promoting immune evasion. Our identification of the specific functional isoform of circTMTC3 may stimulate further investigation into the dynamics and role of circRNA isoforms in immuno-oncology.

In the recent study by Sun *et al*. [75], it was reported that circMGA, a circRNA with tumor-suppressing properties, functions as a chemoattractant for CD8+ T cells. The study indicated a positive correlation between elevated levels of circMGA and improved response to immunotherapy in bladder cancer. However, the analysis conducted by Sun *et al*. solely encompassed a small cohort of 73 patients, without considering the effects of checkpoint blockade immunotherapy. To validate their findings and assess the clinical relevance of circMGA, we performed an expanded analysis employing the Bellmunt *et al*. BLCA cohort [49] (IMvigor010, NCT02450331), which comprised 728 clinical patients encompassing two treatment modalities (374 patients received adjuvant immunotherapy, while 354 were assigned to observation arm). Our analysis demonstrated that heightened levels of circMGA were associated with a favorable prognosis in patients, regardless of their receipt of immunotherapy (Supplementary Figure 2A-B). Notably, we observed that increased circMGA expression presented as a particularly strong predictor of improved overall survival (OS) in early-stage tumors (≤ pT2) received immunotherapy (HR_low vs high_ = 3.77, 95% CI: 1.43-9.94, *P* = 0.0042) (Supplementary Figure 2A). Furthermore, patients with elevated circMGA expression in the immunotherapy group exhibited a higher CD8 T cell signature (Supplementary Figure 2C), potentially clarifying the augmented effectiveness of immunotherapy in these individuals. These findings provide additional substantiation for the conclusions drawn by Sun *et al*.

Together, the two aforementioned case studies, employing analysis of large-scale clinical cancer cohorts, have unveiled distinct impacts and clinical implications of different circRNA expression in cancer immunotherapy across cancer types. These studies showcase the exploratory and validating value of TCCIA in investigating the role of circRNA in tumor immunity.

## Discussion

Circular RNAs, emerging as pivotal gene expression regulators, exert diverse functions across biological processes, with substantial clinical research potential [76]. However, the existing landscape of circRNA-related tumor-immune checkpoint research falls short of satisfying the burgeoning need for insights. This limitation is compounded by constraints stemming from sample quantity and diversity, which subsequently curtail the generalizability of research findings due to geographical, racial, and tumor-specific factors. Consequently, given this evolving landscape, the urgency and significance of developing a CircRNA tool for preliminary data mining have become increasingly pronounced. TCCIA (the Cancer CircRNA Immunome Atlas) an integrated online platform, building upon datasets from four genome sequence archive repositories, encompassing around 4,000 cancer samples, spanning five cancer types and 25 checkpoint-blockade immunotherapy cohorts, incorporating over 0.55 million circRNA expression profiles, 255 established cancer signatures, and TME decomposition results from eight immune infiltration algorithms. TCCIA emphasizes user-friendly visual presentations, eliminating the need for intricate programming skills. Moreover, the platform offers a range of customizable visualization options, ensuring adaptability to user needs. Notably, TCCIA is readily accessible without mandatory registration or login, potentially rendering it an economical and efficient solution for both researchers and clinical practitioners.

In this manuscript, we delineate the data sources, collection and standardization processes, TCCIA’s functionalities, website analysis modules, and provide a step-by-step guide to its operation. To further illustrate TCCIA, we present two concrete examples of the circTMTC3 and circMGA as molecular markers in melanoma and bladder cancer, respectively.

While TCCIA has various advantages and uniqueness, there are still some limitations. In terms of data, although TCCIA includes data from 10 types of cancer, the sample size for some cancer types (e.g., head and neck cancer) remains sparse. Recently, the application of immunotherapy has been expanding to more cancer types, such as colorectal cancer. The main reason for this is that current clinical genomics research of such cancer types primarily focuses on the DNA level, specifically on whole exome sequencing and targeted sequencing [77]. As a result, there are few RNA-seq datasets to infer the presence of circRNAs. Another reason is that some raw RNA-seq datasets are difficult to access due to restricted availability, e.g., Carroll *et al*. study [78]. In terms of data integration, we acknowledge that while TCCIA supports the integration and display of multiple cohorts, there are inherent challenges when it comes to providing uniform functionality for multi-cohort analyses. These challenges arise from the heterogeneity in circRNA profiles and phenotypic data across different cohorts. For users with specific, detailed analysis needs that span multiple cohorts, we recommend leveraging the flexibility of the platform to perform combined analyses based on the results obtained from single-cohort analyses. This approach allows for a more tailored and comprehensive exploration of individual cohort data while accommodating the inherent variations in circRNA profiles and phenotypes across different cohorts. In terms of functionality, some databases provide the characteristics of studying circRNA-miRNA-gene regulatory networks (such as CSCD2 [34], circMine [38], and CircNet2 [35]), which have not been considered in TCCIA with two reasons: First, further experimental validation is generally required to confirm the authenticity of detected circRNAs. In the recent large-scale circRNA detection benchmark study [79], Vromman *et al*. recommended using qPCR+ Ribonuclease R or qPCR+amplicon sequencing for circRNA validation. Therefore, it is recommended to incorporate these experimental validation methods to ensure the accuracy of circRNA detection. Second, the focus of our study is on integrating circRNA profiles and clinical outcomes in cancer patients treated with immunotherapy. However, instead of duplicating efforts, users can leverage already well-established and high-quality circRNA databases to complete other circRNA annotation and analysis explorations. By linking to these databases (a widget is provided in the footer of the TCCIA website), users can access comprehensive circRNA information and utilize existing tools for further analysis, optimizing the accuracy and efficiency of circRNA annotation and deep investigation.

Looking ahead, we aim to continually update TCCIA by incorporating more circRNA data from diverse and updating cancer immunotherapy cohorts and introducing new functionalities based on user feedback. In summary, the distinctive features, analytical capabilities, and potential for growth position it as a pivotal tool in advancing our understanding of circRNAs in tumor immunity and in shaping development of personalized immunotherapy strategies guided by circRNA.

## Methods

### Data collection

To conduct a systematic search, we utilized PubMed (https://www.ncbi.nlm.nih.gov/pubmed/) to search for articles related to Bulk RNA-seq data from solid cancer patients treated with immune checkpoint blockers (ICB). The search expression used was "(ICB [Title/Abstract] OR PD-1[Title/Abstract] OR PD-L1[Title/Abstract] OR CTLA-4[Title/Abstract]) AND (rnaseq[Title/Abstract] OR rna-seq[Title/Abstract] OR rna-sequencing[Title/Abstract] OR rna seq[Title/Abstract] OR rna sequencing[Title/Abstract])". No filters were applied, and there were no restrictions on language or geographic region. Peer-reviewed publications, preprints, and press releases were considered for inclusion. To obtain raw RNA sequencing data, we submitted requests to the Database of Genotypes and Phenotypes (dbGaP) (https://dbgap.ncbi.nlm.nih.gov/), the European Genome-phenome Archive (EGA) (https://ega-archive.org/), and the Genome Sequence Archive (GSA) (https://ngdc.cncb.ac.cn/gsa/) of the National Genomics Data Center (NGDC) after receiving approval from the Data Access Committee (DAC). However, it is important to note that not all raw RNA-seq datasets were accessible and available for use. In total, we collected 23 studies [48–70] related to checkpoint immunotherapy that provided raw RNA-seq datasets (Supplementary Table 1). We gathered relevant clinical data from publications and clinical meta documents associated with these RNA-seq datasets. Additionally, we extracted information on the cohorts’ fundamental characteristics, such as sample size, treatment methods, and specific drugs used, based on the abstracts. It should be noted that, for Patil *et al*. study, two clinical cohorts were included; and for the study identified as PHS000452, the two patient subgroups had distinct drug treatments and clinical annotations. Hence, we treated them as two separate cohorts during the analysis. For the TCCIA project, CircRNA profiles, immunotherapy response, and clinical benefits were analyzed for 10 cancer types. This analysis included over 4,000 clinical samples from 25 cohorts treated with immune-checkpoint blockades (ICBs) such as PD-1/PD-L1 and CTLA-4 inhibitors alone or in combination with conventional therapeutics. The analysis considered both pre-treatment and on-treatment responses.

### CircRNA identification and differential expression analysis

We employed multiple circRNA detection tools, including CIRCexplorer2 [71], CIRIquant [72], find_circ [73], and circRNA_finder [74], to identify, parse, and annotate circRNA junctions within each sample, using the human genome version hg38 as the reference. In our pursuit of biologically robust circRNA identification within each cohort, we undertook an approach that amalgamated results from various circRNA detection tools and implemented a rigorous ensemble strategy characterized by the following stringent criteria. Firstly, we exclusively retained circRNAs located on chromosomes 1 to 22, as well as the X and Y chromosomes. Secondly, circRNAs were mandated to exhibit genomic overlap with gene regions as defined in the reference genome annotation file, specifically "gencode.v34.annotation.gtf." Lastly, to ensure heightened robustness, circRNAs had to be concurrently identified by a minimum of two out of the four employed detection tools, with at least one of these tools detecting no fewer than two back-splicing junction counts. This meticulous methodological approach was instrumental in ensuring the acquisition of biologically sound and reliable circRNA datasets, thereby fortifying the integrity of our analysis within each cohort. Subsequently, the back-splicing junction counts underwent differential expression analysis using established approaches, namely Limma Voom [80], edgeR [81], and DESeq2 [82]. This analysis facilitated comparisons between patient groups, such as those who responded to checkpoint immunotherapy (responders) versus non-responders, as well as between samples collected before and during ICB treatment.

### TME decomposition and cancer gene signature estimation

We employed IOBR [83] for TME decomposition and the scoring of cancer gene signatures. IOBR seamlessly integrates eight widely-used open-source deconvolution methods, including CIBERSORT [84], ESTIMATE [85], quanTIseq [86], TIMER [87], IPS [37], MCPCounter [88], xCell [89], and EPIC [90]. Furthermore, IOBR incorporates a comprehensive compilation of 255 established cancer signatures. These diverse signatures are organized into three distinct categories: TME-associated, tumor-metabolism, and tumor-intrinsic signatures.

### TCCIA implementation

The TCCIA database is developed as a Web application leveraging R Shiny (https://shiny.posit.co/) and built using the golem framework (https://github.com/ThinkR-open/golem) to achieve optimization. TCCIA, is developed solely for research purposes and does not utilize any cookies or collect any personal identifiable information. We independently deployed TCCIA on both the reliable hiplot cloud platform (https://shiny.hiplot.cn/TCCIA/) and our team’s servers (http://tccia.zmu-zhoulab.com/ or http://biotrainee.vip:18888/TCCIA/) for free access. We’re dedicated to promptly addressing and resolving any issues impacting user experience. We will continually expand the TCCIA database, incorporate additional cancer types, and strive to gain access to more data resources every six months.

### Statistical analysis

We performed Kaplan-Meier survival analysis to generate and compare survival curves. The log-rank test was used for comparison. We also conducted multivariate survival analysis using the Cox regression model. All reported *P*-values are two-tailed, and a significance level of p≤0.05 was used unless otherwise specified. All statistical analyses and visualization were conducted using R v4.2.0.

### Patient consent for publication

Not applicable.

### Ethics

In this study, we conducted a secondary analysis of trial datasets, which was determined to carry minimal risk. The Institutional Review Board of the Second Affiliated Hospital, Zunyi Medical University (No. YXLL(KY-R)-2021-010) approved the study protocol. As per national legislation and institutional guidelines, written informed consent for participation was not deemed necessary for this particular study.

### Data availability

All the processed data can be accessed on the TCCIA website or in the article. We collected ICB-related cancer data sets with tumor gene expression profiles from the dbGaP, EGA, EMBL-EBI and GSA databases. Following the accession instruction described in published ICB studies (Table S1), we downloaded ICB patients’ RNA-Seq raw sequencing data, clinical information, and response outcome information from ICB studies (if available). Curated circRNA identification results from 4 detection methods and our ensemble approach are available at Zenodo (https://zenodo.org/doi/10.5281/zenodo.7969298). For requesting processed circRNA expression data from the TCCIA, please contact the project leader, Jian-Guo Zhou.

### Code availability

The cancer circRNA identification pipeline with an ensemble approach and corresponding scripts and logs for handling TCCIA cohorts are available at https://github.com/ShixiangWang/circrna-pipeline (commit id: 1115905).

### Funding

This work was supported by the National Natural Science Foundation of China (Grant No. 81660512, 81472594, 81770781, 82270825), Chunhui program of the Chinese Ministry of Education (Grant No. HZKY20220231), the Natural Science Foundation of Guizhou Province (Grant No. ZK2021-YB435), Guangdong Basic and Applied Basic Research Foundation (Grant No. 2021A1515011743), Youth Talent Project of Guizhou Provincial Department of Education (Grant No. QJJ2022-224), China Postdoctoral Science Foundation (Grant No. 2021M703733), China Lung Cancer Immunotherapy Research Project, Special funds for innovation in Hunan Province (Grant No. 2020SK2062), the Science and Technology Programs of Zunyi City (Grant No. ZSKH-HZ2023-142), and Collaborative Innovation Center of Chinese Ministry of Education (Grant No. 2020-39).

### Contributions

SW: Conceptualization, software, methodology, formal analysis, writing original draft, review and editing. YX: Software, methodology, formal analysis, visualization, writing original draft. YZ: Software, formal analysis, visualization, writing original draft. HW: Conceptualization, writing original draft, review and editing. JL and USG: Resources, review and editing. MC: Visualization, review and editing. PL, YHL, MH and BF: Review and editing. XL, QZ and HM: Supervision, resources, funding acquisition. JGZ: Conceptualization, methodology, resources, supervision, funding acquisition, project administration, writing original draft, writing–review and editing.

### Conflict of interest

None were declared.

## Acknowledgments

We thank Dr. Jianming Zeng (University of Macau), and all the members of his bioinformatics team, biotrainee, for generously sharing their experience and codes. We thank Juan Zhang (Echo Biotech Co., Ltd., Beijing, China) for her help in the data pre-processing. We thank Jieyin Tan (University of Sydney) for test TCCIA and report bugs. The present work was performed in (partial) fulfillment of the requirements for obtaining the degree “Dr. rer. biol. hum.”at the Friedrich-Alexander-Universität Erlangen-Nürnberg (FAU).

---

**Supplementary Table1. TCCIA cohort summary.**

**Abbr.** BLCA, Bladder urothelial carcinoma; HNSC, Head and neck squamous cell carcinoma; KIRC, Kidney renal clear cell carcinoma; NSCLC, Non-small lung cancer; SCLC, Small lung cancer; SKCM, Skin cutaneous melanoma; SGC, Salivary gland carcinoma; BRCA, Breast cancer; SARC, Sarcoma; STAD, Stomach Adenocarcinoma.

**Supplementary Figure 1. CircRNA identification and overlap analysis of TCCIA cohorts.** (**A**)The number of detected circRNAs in each cohort by different method. (**B**) t-SNE plot of all samples, colored by cohorts.

**Abbr.** BLCA, Bladder urothelial carcinoma; HNSC, Head and neck squamous cell carcinoma; KIRC, Kidney renal clear cell carcinoma; NSCLC, Non-small lung cancer; SCLC, Small lung cancer; SKCM, Skin cutaneous melanoma; SGC, Salivary gland carcinoma; BRCA, Breast cancer; SARC, Sarcoma; STAD, Stomach Adenocarcinoma.

**Supplementary Figure 2. circMGA as a predictive biomarker for adjuvant immunotherapy in bladder cancer.** (**A**) Overall survival (OS) of bladder cancer patients (all patients, pathological tumor stage ≤pT2, pT3/4) treated with adjuvant immunotherapy was analyzed by Kaplan–Meier curves. Red, group of high circRNA expression. Blue, group of low circRNA expression. We used the maximally selected rank statistics method to determine the cutoff. The minimal proportion for optimal cutoff was set as 0.4. P value was calculated using a log-rank test. (B) Overall survival (OS) of bladder cancer patients (all patients, early-stage tumors(≤ pT2) and advanced-stage tumors (pT3/4)) treated with adjuvant immunotherapy was analyzed by Kaplan–Meier curves. Red, group of high circRNA expression. Blue, group of low circRNA expression. We used the maximally selected rank statistics method to determine the cutoff. The minimal proportion for optimal cutoff was set as 0.4. P value was calculated using a log-rank test. (C) Boxplot visualizing the association of circMGA subgroups for patients receiving adjuvant immunotherapy with CD8 T cells signature.

## Notes

http://tccia.zmu-zhoulab.com/

http://biotrainee.vip:18888/TCCIA/

https://shiny.hiplot.cn/TCCIA

### Competing Interest Statement

The authors have declared no competing interest.

### Summary of Updates

R1

